# A meiotic drive element in the maize pathogen *Fusarium verticillioides* is located within a 102-kb region of chromosome V

**DOI:** 10.1101/045914

**Authors:** Jay Pyle, Tejas Patel, Brianna Merril, Chabu Nsokoshi, Morgan McCall, Robert H. Proctor, Daren W. Brown, Thomas M. Hammond

## Abstract

*Fusarium verticillioides* is an agriculturally important fungus because of its association with maize and its propensity to contaminate grain with toxic compounds. Some isolates of the fungus harbor a meiotic drive element known as Spore *killer* (*Sk*^*K*^) that causes nearly all surviving meiotic progeny from an *Sk*^*K*^ × Spore killer-susceptible (*Sk*^*S*^) cross to inherit the *Sk*^*K*^ allele. *Sk*^*K*^ has been mapped to chromosome V but the genetic element responsible for meiotic drive and spore killing has yet to be identified. In this study, we used cleaved amplified polymorphic sequence markers to genotype individual progeny from an *Sk*^*K*^ × *Sk*^*S*^ mapping population. We also sequenced the genomes of three progeny from the mapping population to determine their single nucleotide polymorphisms. These techniques allowed us to refine the location of *Sk*^*K*^ to a contiguous 102-kb region of chromosome V, herein referred to as the *Sk* locus. Relative to *Sk*^*S*^ genotypes, *Sk*^*K*^ genotypes have one extra gene within this locus for a total of 42 genes. The additional gene in *Sk*^*K*^ genotypes, named SKL1 for *Spore Killer Locus 1*, is the most highly expressed gene from the *Sk* locus during early stages of sexual development. The *Sk* locus also has three hypervariable regions, the longest of which includes *SKL1*. The possibility that *SKL1*, or another gene from the *Sk* locus, is an essential component of meiotic drive and spore killing is discussed.

## INTRODUCTION

*Fusarium verticillioides* is an ascomycete fungus that can exhibit both endophytic and pathogenic growth on maize (Bacon and Hinton 1996; White 1998). The fungus is of agriculture concern because of its ability to cause maize ear and stalk rot, and also because it can contaminate maize kernels with a group of mycotoxins known as fumonisins (Marasas 2001; Brown and Proctor 2016). The problems caused by *F. verticillioides* and its myoctoxins were noticed as early as 1970, when the fungus was correlated with an outbreak of Equine Leukoencephalomalacia (ELEM) in South Africa (Kellerman *et al.* 1972). Since then, fumonisins have been confirmed to cause ELEM in horses (Marasas *et al.* 1988; Kellerman *et al.* 1990), pulmonary edema in pigs (Harrison *et al.* 1990), and liver cancer in laboratory rats (Gelderblom *et al.* 1988, 1991). Fumonisin-contaminated grain has also been linked to neural tube defects in humans (Missmer *et al.* 2006). Thus, the potential presence of fumonisins, even in apparently healthy grain, makes it necessary to aggressively screen grain for these mycotoxins before it is used in food or feed. Contaminated grain must be destroyed, resulting in crop losses worth millions of dollars per year (Wu, 2007).

The primary sources of inoculum for endophytic and pathogenic growth of *F. verticillioides* on maize are asexual spores, such as macroconidia and microconidia (Munkvold 2003). Sexual spores, called ascospores, may also be important to the *F. verticillioides* life cycle (Reynoso *et al.* 2009). Sexual reproduction in this heterothallic fungus begins when the immature fruiting bodies of one strain are fertilized by a strain of opposite mating type (Martin *et al.* 2011). Fruiting bodies are called perithecia and meiosis occurs in spore sacs, called asci, within the perithecia. At the start of meiosis, a single diploid nucleus is formed from the haploid genomes of both parents. This diploid nucleus separates into four haploid nuclei through the standard reductional and equational division stages of meiosis, and subsequently, each of these four haploid nuclei undergo a post-meiotic mitosis to form eight nuclei. Each nucleus is then incorporated into a separate ascospore through a process known as ascospore delimitation. A phenotypically normal ascus should thus contain eight ascospores at maturity in *F. verticillioides*.

During normal development, asci break down and ascospores are exuded from perithecia in gelatinous masses. However, it is possible to examine ascospores within intact asci by perithecial dissection. Kathariou and Spieth used this procedure to discover an allele named Spore killer (*Sk*^*K*^). In *Sk*^*K*^ × *Sk*^*S*^ crosses, where the latter stands for *Spore killer*-susceptible, asci contain four instead of eight ascospores (Kathariou and Spieth 1982). Early stages of ascus development are cytologically-normal in these crosses, but four ascospores degenerate shortly after ascospore delimitation (Raju 1994). The four surviving ascospores are almost always of the *Sk*^*K*^ genotype (Kathariou and Spieth 1982; Xu and Leslie 1996). Unlike *Sk*^*K*^ × *Sk*^*S*^ crosses, *Sk*^*K*^ × *Sk*^*K*^ crosses produce asci with eight ascospores (Kathariou and Spieth 1982). Thus, the *Sk*^*K*^ allele is sufficient for spore killing and resistance to spore killing.

The ability of *Sk*^*K*^-ascospores to kill *Sk*^*S*^-ascospores suggests that *Sk*^*K*^ could be the predominant allele in some *F. verticillioides* populations. Although the literature is limited on this subject, a screen of 225 *F. verticillioides* isolates from 24 fields across Europe and North America found the *Sk*^*K*^ allele to be present in 81% of isolates (Kathariou and Spieth 1982). Shortly after the discovery of *Sk*^*K*^ in *F. verticillioides*, a spore killing allele was also discovered in *F. subglutinans.* This allele was also named Spore killer, but it was given the slightly different notation of *SK*^*k*^. Along with the report of its discovery, Sidhu (1984) presented evidence that the *SK*^*k*^ allele was present in 10 of 15 F. subglutinans isolates obtained from a single maize field. Therefore, *Sk*^*K*^ and *SK*^*k*^ appear to be the predominant alleles in at least some populations of *F. verticillioides* and *F. subglutinans.*

The relationship between *Sk*^*K*^ and *SK*^*k*^ is still unclear despite both being discovered more than 30 years ago. To our knowledge, the only other significant research involving either allele was performed by Xu and Leslie (1996) during construction of a genetic map for *F. verticillioides*. During this study, *Sk*^*K*^ was mapped between two restriction fragment length polymorphism (RFLP) markers on chromosome V. These markers, named RFLP1 and 11p18, are located 2.5 cM and 8.6 cM from *Sk*^*K*^, respectively (Xu and Leslie 1996). Xu and Leslie's mapping population was derived from an *Sk*^*K*^ × *Sk*^*S*^ cross, and interestingly, only one progeny from over 100 did not inherit the *Sk*^*K*^ allele (Xu and Leslie 1996). This evidence demonstrates that *Sk*^*K*^ can achieve transmission rates of over 99 percent in laboratory crosses.

An *F. verticillioides* reference genome, derived from an *Sk*^*S*^ strain known as Fv149-*Sk*^*S*^, was published 14 years after the initial mapping of *Sk*^*K*^ (Ma *et al.* 2010). Here, we advance Fusarium Spore killer research by delineating physical borders for *Sk*^*K*^ with respect to the *F. verticillioides* reference genome. Our data place *Sk*^*K*^ within a 102-kb contiguous sequence of DNA, which we refer to as the *Sk* locus. Notable differences exist between the *Sk* locus in Fv999-*Sk*^*K*^ and Fv149-*Sk*^*S*^ strains. These are described and discussed below.

## MATERIAL AND METHODS

### Strains, media and culture conditions

Key strains are listed in Table 1. Vegetative propagation was performed on V8 Juice Agar (VJA) (Tuite, 1969) in 16 mm test tubes at room temperature on a laboratory benchtop. Carrot agar (CA), which was originally described by Klittich and Leslie (1988), was prepared as follows: 200 g of organic peeled baby cut carrots were autoclaved in 200 ml of water, pureed with a blender, then adjusted to 500 ml with sterile water to create a 1× stock. Diluted CA (e.g., 0.1× and 0.25×) was prepared by mixing the appropriate volumes of 1× CA stock and sterile water, before adding agar to a final concentration of 2% and autoclaving. Liquid Vogel’s Minimal Medium (VMM, Vogel 1956) or GYP medium (2% glucose, 1% peptone, and 0.3% yeast extract) were used to produce mycelia for genomic DNA isolation. Liquid cultures for genomic DNA isolation were incubated at 28 °C in the dark without agitation.

**Table 1.**
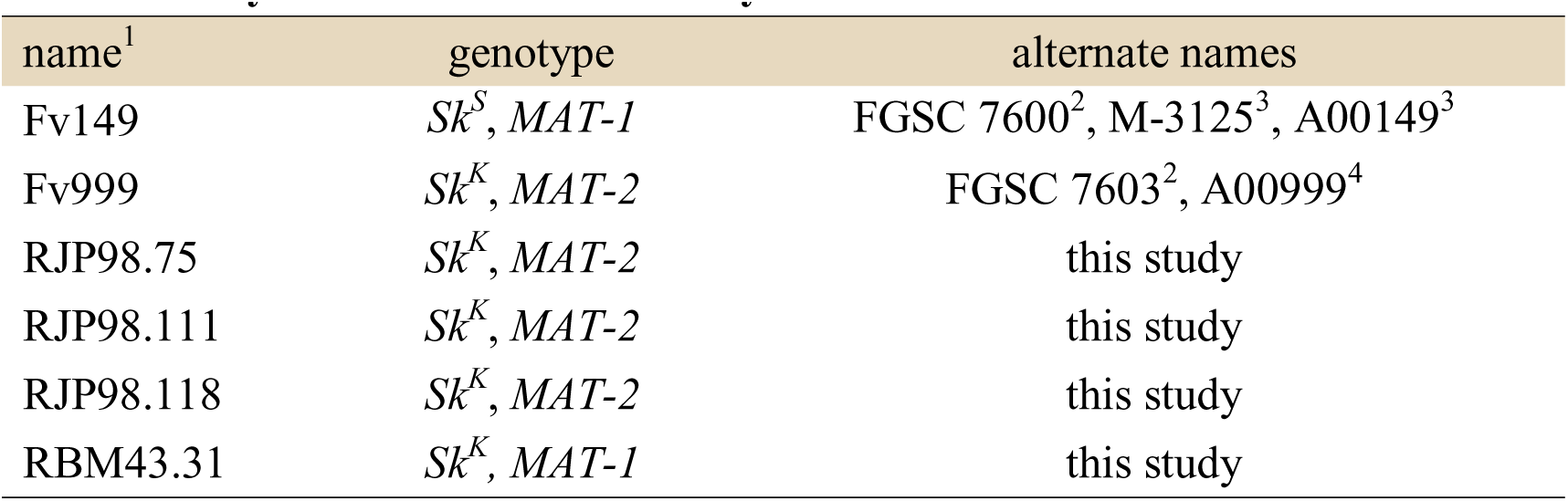
Key strains used in this study. ^1^Throughout the text, the suffix -*Sk*^*K*^ or -*Sk*^*S*^ is added to each strain name to denote genotype with respect to Sk. ^2^McCluskey *et al.* 2010; ^3^Xu *et al.* 1995; ^4^Xu and Leslie 1996.

| name<sup>1</sup> | genotype | alternate names |
| --- | --- | --- |
| Fv149 | $Sk^S$, *MAT-1* | FGSC 7600<sup>2</sup>, M-3125<sup>3</sup>, A00149<sup>3</sup> |
| Fv999 | $Sk^K$, *MAT-2* | FGSC 7603<sup>2</sup>, A00999<sup>4</sup> |
| RJP98.75 | $Sk^K$, *MAT-2* | this study |
| RJP98.111 | $Sk^K$, *MAT-2* | this study |
| RJP98.118 | $Sk^K$, *MAT-2* | this study |
| RBM43.31 | $Sk^K$, *MAT-1* | this study |

### Sexual crosses

Crosses were performed on CA in a similar manner to previously described methods (Klittich and Leslie 1988). Asexual spores (conidia) from the female parent were transferred to the center of a 60 mm petri dish containing 20 ml of CA or diluted CA. The female parent was then cultured for 10-14 days before fertilization with a suspension of conidia from the male parent. Conidial suspensions were prepared by adding 2.0 ml of 0.001% Tween-20 to a 10-14 day old test-tube culture of the male parent and dislodging the conidia with a pipette tip. Fertilization was performed by transferring 1.0 ml of a conidial suspension of the male parent to the surface of a culture of the female parent. The conidial suspension was spread over the surface of the female culture with a glass rod. Crosses were incubated in a culture chamber that alternated between 23.0 °C (12 hours, light) and 22.5 °C (12 hours, dark). Light was provided by white (Philips F34T12/CW/RS/EW/ALTO) and black (General Electric F40T12BL) fluorescent lamps.

### The Fv999-*Sk*^*K*^ × Fv149-*Sk*^*S*^ mapping population

In *F. verticillioides*, ascospores are exuded from mature perithecia in a hair-like structure called a cirrus. To obtain an *Sk*^*K*^ mapping population, Fv999-*Sk*^*K*^ was crossed with Fv149-*Sk*^*S*^ and cirri were isolated from the tops of a few perithecia with a sterile needle, dispersed in sterile water, and spread onto a plate of 4% water agar. Germinating ascospores were transferred to VJA in 16 mm test tubes.

### Microscopy

Asci were dissected from perithecia in 25% glycerol under magnification. A Vanguard 1433Phi light microscope and attached digital camera (Amscope MU1000) were used for imaging. The condenser and aperture diaphragm were set for high contrast, which allowed for the number of ascospores in mature asci to be determined without tissue staining.

### DNA methods

Genomic DNA was prepared using one of three methods. In method one, strains were cultured in 25 ml liquid VMM at 28 °C in the dark for three days. Mycelia were washed with 0.9% NaCl and dried by lyophilization before extraction with IBI Scientific’s Genomic DNA Mini Kit for Plants. Method two, based on Henderson *et al.* (2005), was used as an inexpensive and time-efficient alternative to method one. A 6” plain-tipped wood applicator was used to transfer ≤ 10 mg of conidia to 200 μl of TE buffer (10 mM Tris, 1 mM EDTA, pH 8.0). The suspension was boiled at 105 °C in a heat block for 12 minutes, incubated on ice for 2 minutes, then vortexed for 5 seconds. Insoluble material was pelleted at 15,000 × g for 10 minutes at room temperature, after which 25 μl of supernatant was transferred to a new vial and frozen at −20 °C for storage.In our hands, this method works well for the amplification of PCR products shorter than 1 kb. Method three was used for the preparation of genomic DNA for high-throughput sequencing. Strains were cultured in 25 ml of liquid GYP at 28 °C for 48 hrs. Mycelia was washed with water and DNA was extracted with the Zymo Research Fungal DNA Miniprep Kit.

Polymerase chain reaction (PCR) assays were performed with MidSci Bullseye Taq DNA Polymerase or New England Biolabs Phusion High-Fidelity DNA Polymerase.

### Genome sequencing

DNA libraries for MiSeq sequencing (Illumina) were constructed from 1 ng of genomic DNA using the Illumina Nextera XT DNA Library Preparation Kit. Sequencing was performed with MiSeq Reagent Kit Version 3. Adapters were removed and low-quality reads were trimmed with CLC Genomics Workbench (Version 8.0). The datasets were deposited in the National Center for Biotechnology Information’s (NCBI) Sequence Read Archive (Leinonen *et al.* 2011). They can be obtained with the following accession numbers: SRR3271586 (Fv999-*Sk*^*K*^), SRR3273544 (JP98.75-*Sk*^*K*^), SRR3273545 (JP98.111-*Sk*^*K*^), and SRR3273546 (JP98.118-*Sk*^*K*^). The data sets are of high-quality. For example, draft genomes were assembled with CLC Genomics Workbench and all assemblies had N50 values over 90 kb with coverage levels between 49 and 73 fold (Table 2).

**Table 2.** Draft genome assembly statistics. ^1^total, the total length of each genome assembly; ^2^avg. length, the average length of each read in the assembly; ^3^coverage, calculated by multiplying the number of reads by the average read length and dividing the product by 41.9 Mb, the genome size of Fv149-*Sk*^*S*^ (NCBI ASM14955v1).

| name | total<sup>1</sup> | contigs | avg. length<sup>2</sup> | # reads | N50 | coverage<sup>3</sup> |
| --- | --- | --- | --- | --- | --- | --- |
| Fv999-*Sk<sup>K</sup>* | 41,922,529 | 855 | 255 | 10,956,620 | 109,413 | 67 |
| RPJ98.75-*Sk<sup>K</sup>* | 41,848,894 | 985 | 257 | 8,087,199 | 91,461 | 50 |
| RPJ98.111-*Sk<sup>K</sup>* | 41,892,017 | 841 | 242 | 12,538,869 | 120,565 | 72 |
| RPJ98.118-*Sk<sup>K</sup>* | 41,868,129 | 914 | 268 | 9,009,572 | 88,867 | 58 |

### CAPS markers

Cleaved amplified polymorphic sequence (CAPS) markers (Konieczny and Ausubel 1993) were used to help refine the location of *Sk*^*K*^ (Table 3). CAPS markers were identified by first aligning genome sequencing reads from strain Fv999-*Sk*^*K*^ to the Fv149-*Sk*^*S*^ reference genome (NCBI, ASM14955v1) and then visually scanning aligned reads with Tablet (Milne *et al.* 2010) for polymorphisms in GGCC sites. This four-base sequence is cleaved by the restriction endonuclease *Hae*III. Six CAPS markers on chromosome V were chosen for this study, along with one CAPS marker on each of chromosomes I, VII, and XI (Table 3). PCR-primers for each CAPS marker were designed to amplify a short product (< 500 bp) from both Fv999-*Sk*^*K*^ and Fv149-*Sk*^*S*^ sequences. The PCR primers used for amplification of each CAPS marker are described in Table S1.

**Table 3.** Marker locations. Marker locations on chromosomes I, V, VII, and XI of *F. verticillioides* Fv149-*Sk*^*S*^ (NCBI ASM14955v1).

| marker | chromosome | position (Mb) |
| --- | --- | --- |
| CAPS-1 | V | 1.47 |
| CAPS-2 | V | 1.42 |
| CAPS-3 | V | 0.78 |
| CAPS-4 | V | 0.43 |
| CAPS-5 | V | 0.02 |
| CAPS-6 | V | 2.12 |
| CAPS-9 | XI | 0.07 |
| CAPS-10 | VII | 1.05 |
| CAPS-11 | I | 0.73 |
| RFLP-11p18 | V | 1.67 |

The following protocol was used to analyze the segregation patterns of CAPS markers in each individual of the Fv999-*Sk*^*K*^ × Fv149-*Sk*^*S*^ mapping population. Genomic DNA was isolated from each progeny, CAPS markers were amplified by PCR, and PCR products were digested with *Hae*III. The digested products were then examined for Fv999-*Sk*^*K*^ or Fv149-*Sk*^*S*^ cleavage patterns by gel electrophoresis on standard 1% agarose-TAE (40mM Tris, 20mM acetic acid, and 1mM EDTA) gels. The *Hae*III-based DNA digest patterns of each CAPS marker for Fv999-*Sk*^*K*^ and Fv149-*Sk*^*S*^ alleles are listed in Table S2.

### Single nucleotide polymorphisms (SNPs)

Reads from each MiSeq dataset were aligned to the reference genome of strain Fv149-*Sk*^*S*^ with Bowtie 2 (Langmead and Salzberg 2012). SAMtools 1.3 and BCFtools 1.3 were then used to report SNPs in variant call format (VCF) (Li *et al.* 2009; Danecek *et al.* 2011). Only reads that matched to a single region of Fv149-*Sk*^*S*^ with less than 10 mismatches were used to produce the VCF files. Custom Perl scripts were then used to extract significant SNPs from the VCF files. Significant SNPs were defined as those which were supported by at least 90% of reads at each position. Only positions covered by more than nine reads were considered. Polymorphisms caused by insertions or deletions were ignored.

### Gene predictions

Protein-coding genes within the *Sk* locus of Fv999-*Sk*^*K*^ were predicted as follows: first, the sequence of the *Sk* locus in Fv999-*Sk*^*K*^ was obtained by de novo assembly of MiSeq reads (Table 2); second, the coding sequences of annotated genes from the *Sk* locus of Fv149-*Sk*^*S*^ were obtained from GenBank (CM000582.1); third, the Fv149-*Sk*^*S*^ coding sequences were aligned to the *Sk* locus of Fv999-*Sk*^*K*^ with Clustal Omega (Sievers *et al.* 2011; Li *et al.* 2015); and fourth, the alignments were used to manually annotate protein-coding sequences within the sequence of the Fv999-*Sk*^*K*^ *Sk* locus. These steps identified 41 putative protein-coding genes within the *Sk* locus of Fv999-*Sk*^*K*^. Augustus 3.2.1 (Stanke and Waack 2003) was then used for *de novo* prediction of protein-coding genes. Augustus predictions matched our manual annotation with the notable exception of an additional identified gene (*SKL1*, described below) within the *Sk* locus of Fv999-*Sk*^*K*^. The complete sequence and annotation of the 102-kb *Sk*^*K*^ locus from Fv999-*Sk*^*K*^ can be obtained from GenBank with accession number KU963213.

## RESULTS

### Increased production of fruiting bodies by strain Fv999-*Sk*^*K*^ on diluted CA

Directional crosses are often used to investigate sexual processes in heterothallic ascomycete fungi. In this type of cross, one parent is designated as the female and the other is designated as the male. The female provides protoperithecia, i.e., immature fruiting bodies, which are fertilized by conidia from the male. After fertilization, protoperithecia develop into perithecia.

Carrot Agar (CA) is commonly used as a growth medium to study sexual processes in *F. verticillioides*. Its preparation essentially involves purchasing carrots from a local market, followed by peeling, blending, and autoclaving a predetermined weight of carrots within a specified volume of water (Leslie and Summerell 2006). Because we were curious about how the amount of carrot in CA medium influences the productivity of Fv999-*Sk*^*K*^ × Fv149-*Sk*^*S*^ crosses, Fv999-*Sk*^*K*^ was cultured on 0.001×, 0.01×, 0.1×, 0.25×, 0.5×, and 1.0× CA, then fertilized with Fv149-*Sk*^*S*^ conidia. The crosses performed on 0.1× and 0.25× CA resulted in nearly three to six fold more perithecia than the crosses performed on 0.5× and 1.0× CA (Figure 1). No perithecia were produced when crosses were performed on 0.001× and 0.01× CA (Figure 1). Additionally, less conidia were produced on 0.1× CA than on 0.25× CA (data not shown). Because conidia can be a source of cross-contamination, 0.1× CA was used as the crossing medium for the remainder of this study.

**Figure 1.**
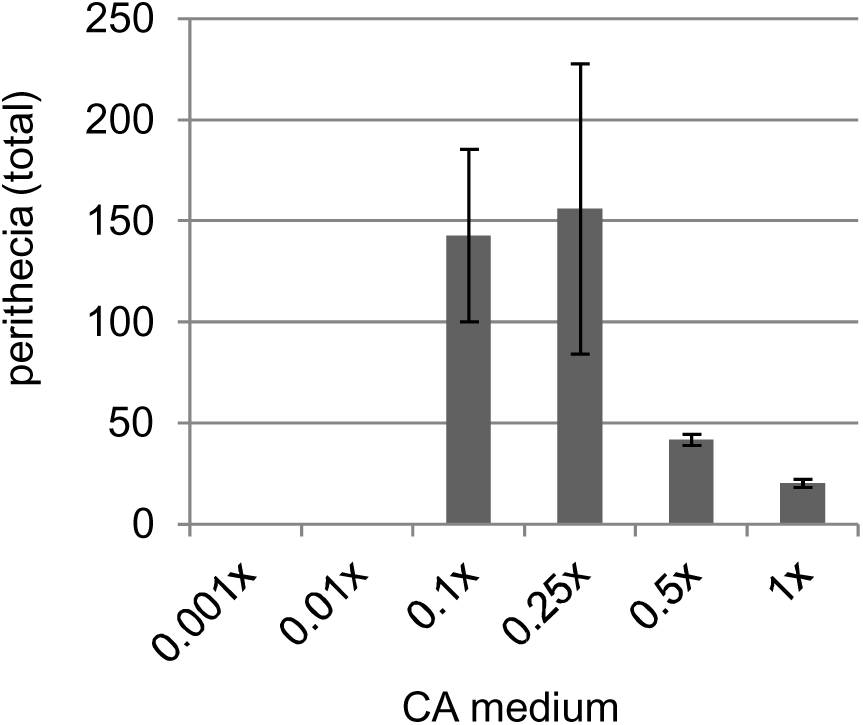
Perithecial production on different CA media. Cultures of Fv999-*Sk*^*K*^ were grown on CA media containing different concentrations of carrot and fertilized with conidia from strain Fv149-*Sk*^*S*^. The data represent the total number of perithecia observed 25 days after fertilization. Crosses were performed in triplicate and error bars are standard deviation.n

### RFLP 11p18 overlaps with FVEG_02851 on chromosome V

Although the sequence of RFLP1 was not available, we obtained the sequence of RFLP 11p18 from the *Fusarium* Comparative Database. We used this sequence as the query in a BLASTn (Altschul *et al.* 1990) search of the *F. verticillioides* reference genome. This search identified positions 1672753 to 1673602 on chromosome V (CM000582.1) as a match to 11p18. This region overlaps with most of the predicted coding sequence of gene *FVEG_02851*, which spans positions 1672886 to 1673769 and encodes a protein of unknown function. Its position on chromosome V was used to choose CAPS markers near the *Sk* locus (Figure 2A).

**Figure 2.**
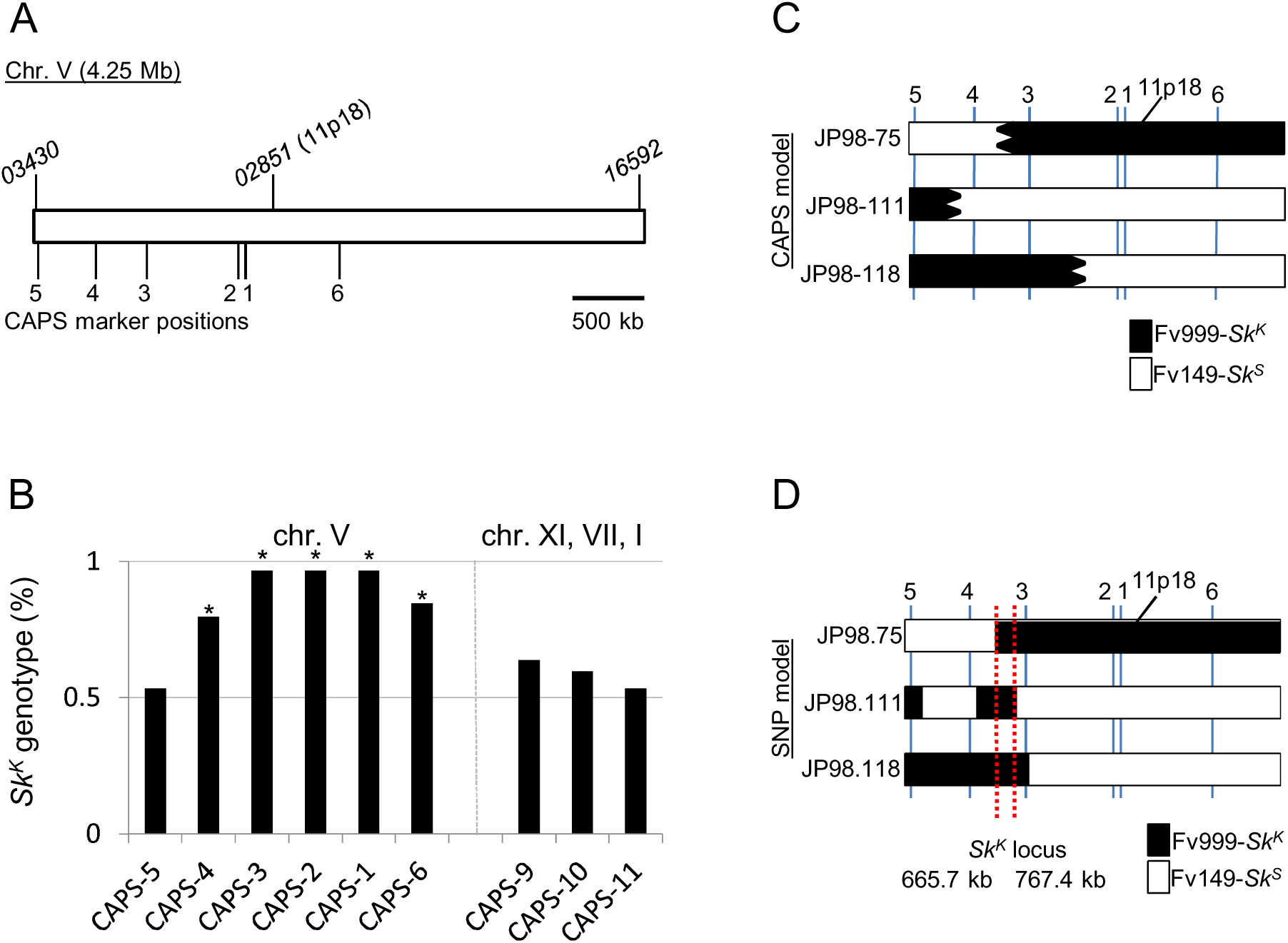
The *F. verticillioides Sk* locus is found within a 102-kb region of chromosome V. A)A diagram of the 4.25 Mb sequence of chromosome V from Fv149-*Sk*^*S*^ is shown. The annotated chromosome V sequence was downloaded from NCBI (CM000582.1). The first gene (*FVEG_03430*) and last gene (*FVEG_16592*) on the chromosome are indicated. The location of RFLP marker 11p18 is also shown. Marker 11p18 overlaps gene *FVEG_02851*. B) A total of 60 progeny were isolated from a cross between Fv999-*Sk*^*K*^ and Fv149-*Sk*^*S*^ These progeny were genotyped with nine different CAPS markers. Markers CAPS-1 through CAPS-6 are located on chromosome V. Markers CAPS-9, CAPS-10, and CAPS-11 are found on chromosomes XI, VII, and I, respectively. Each bar represents the percentage of Fv999-*Sk*^*K*^ genotypes recovered for each marker. An asterisk is placed above all markers whose recovery deviated significantly from a Mendelian ratio of 1:1 according to a χ^2^ test with p × 0.01. The biased transmission of Fv999-*Sk*^*K*^ sequences was detected for all markers on chromosome V, except for CAPS-5. (C) The marker patterns for progeny JP98.75-*Sk*^*K*^, JP98.111-*Sk*^*K*^, and JP98.118-*Sk*^*K*^ suggest that crossover events occurred between CAPS-3 and CAPS-4 (for JP98.75-*Sk*^*K*^), CAPS-4 and CAPS-5 (for JP98.111-*Sk*^*K*^), and CAPS-2 and CAPS-3 (for JP98.118-*Sk*^*K*^) (Table S3). These crossovers do not reveal an Fv999-*Sk*^*K*^-derived chromosome V sequence common to all three progeny. The predicted Fv999-*Sk*^*K*^‐ and Fv149-*Sk*^*S*^-inherited regions are shown in black and white,respectively. Irregular borders are used to indicate that the exact crossover positions cannot be determined from the marker data. (D) SNP analysis was used to more accurately define crossover positions for progeny JP98.75-*Sk*^*K*^, JP98.111-*Sk*^*K*^, and JP98.118-*Sk*^*K*^, and to discover additional crossovers. This analysis identified two additional recombination events on chromosome V of JP98.111-*Sk*^*K*^, which delineate a single contiguous region between 665.7 kb and 767.4 kb on chromosome V as the only Fv999-*Sk*^*K*^ region common to all three progeny (red dotted lines). This region is referred to as the *Sk* locus.

### Loci linked to *Sk*^*K*^ drive through meiosis

To begin refining the location of *Sk*^*K*^ on chromosome V, Fv999-*Sk*^*K*^ was crossed with Fv149-*Sk*^*S*^ and the segregation patterns of nine CAPS markers were examined in 60 progeny. Both Fv999-*Sk*^*K*^ and Fv149-*Sk*^*S*^ patterns were inherited in a 1:1 ratio for chromosomes XI, VII, and I. In contrast, Fv999-*Sk*^*K*^ patterns of CAPS markers on chromosome V were inherited more frequently (>= 79.6%) than expected (Figure 2B and Table S3). The only exception was CAPS-5, which is located approximately 20 kb from a telomere (Figure 2A and Table 3). Overall, these results mark the first independent confirmation of Xu and Leslie’s (1996) original findings on the biased recovery of molecular markers linked to *Sk*^*K*^ in *Sk*^*K*^ × *Sk*^*S*^ crosses.

### Genome sequencing refines the *Sk* locus to a 102-kb region of chromosome V

To delineate the position of *Sk*^*K*^ on chromosome V, we narrowed our focus to progeny JP98.75-*Sk*^*K*^, JP98.111-*Sk*^*K*^, and JP98.118-*Sk*^*K*^. Even though CAPS marker analysis failed to identify an Fv999-*Sk*^*K*^ region of chromosome V that was inherited by all three progeny (Figure 2C, black shaded regions), all were determined to carry the *Sk*^*K*^ allele by phenotypic analysis (Figure 3 and data not shown). Therefore, their genomes were sequenced (Table 2) and their SNPs along chromosome V were examined. This analysis allowed us to identify two additional recombination events. Both of these recombination events occurred between CAPS markers CAPS-3 and CAPS-4 in the ascus producing progeny JP98.111-*Sk*^*K*^ (Figure 2D), explaining why they were not revealed by our CAPS marker analysis. More importantly, the SNP profiles of all three progeny require that *Sk*^*K*^ be located between positions 665,669 and 767,411 (Figure 2D, red lines) with respect to the reference sequence (GenBank, CM000582.1). We refer to this 102-kb contiguous sequence as the *Sk* locus (GenBank, KU963213).

**Figure 3.**
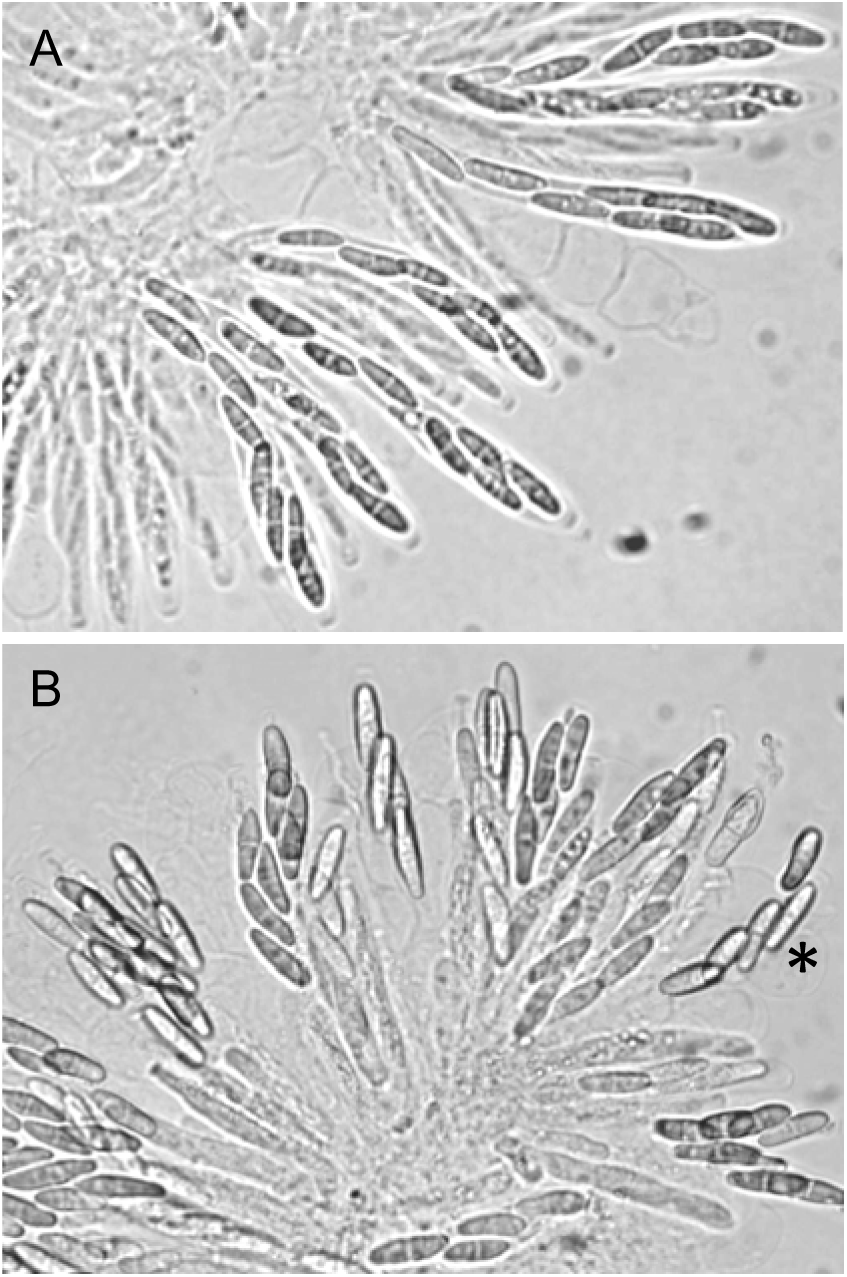
Representative images of killing and no-killing in *F. verticillioides* asci. (A) Asci from a cross of progeny JP98.111-*Sk*^*K*^ with BM43.31-*Sk*^*K*^. Eight ascospores can be detected in most of the mature asci, confirming that progeny JP98.111-*Sk*^*K*^ carries the *Sk*^*K*^ allele. (B) Asci from a cross of progeny JP98.111-*Sk*^*K*^ with Fv149-*Sk*^*S*^. Only four ascospores are detected in most of the mature asci, again confirming that progeny JP98.111-*Sk*^*K*^ carries the *Sk*^*K*^ allele. The asterisk denotes an ascus where there appears to be five ascospores. Two possible explanations are that this 5^th^ ascospore escaped killing, or, it strayed from another ascus during the dissection and imaging process.

### *Sk*^*K*^ strains carry a unique gene in the *Sk* locus

The *Sk* locus spans 102,256 bases in Fv999-*Sk*^*K*^, but only 101,743 bases in Fv149-*Sk*^*S*^. A ClustalW-based alignment (Thompson *et al.* 1994) of the sequences from both strains is 102,557 bases long, with 301 gaps in the Fv999-*Sk*^*K*^ sequence and 814 gaps in the Fv149-*Sk*^*S*^ sequence (Figure 4). To identify specific regions in *Sk*^*K*^ and *Sk*^*S*^ strains with high levels of SNPs or gaps, the number of SNPs or gaps were calculated for each 100-base window across the alignment. A qualitative analysis of these results finds at least three hypervariable regions (Figure 4). One of the shorter hypervariable regions spans genes *FVEG_03180* to *FVEG_03175*, while another spans genes *FVEG_03174* and *FVEG_03173* (Figure 4). The longest hypervariable region is approximately 14 kb long, spans gene FVEG_03167, and extends to the right border of the *Sk* locus (Figure 4).

**Figure 4.**
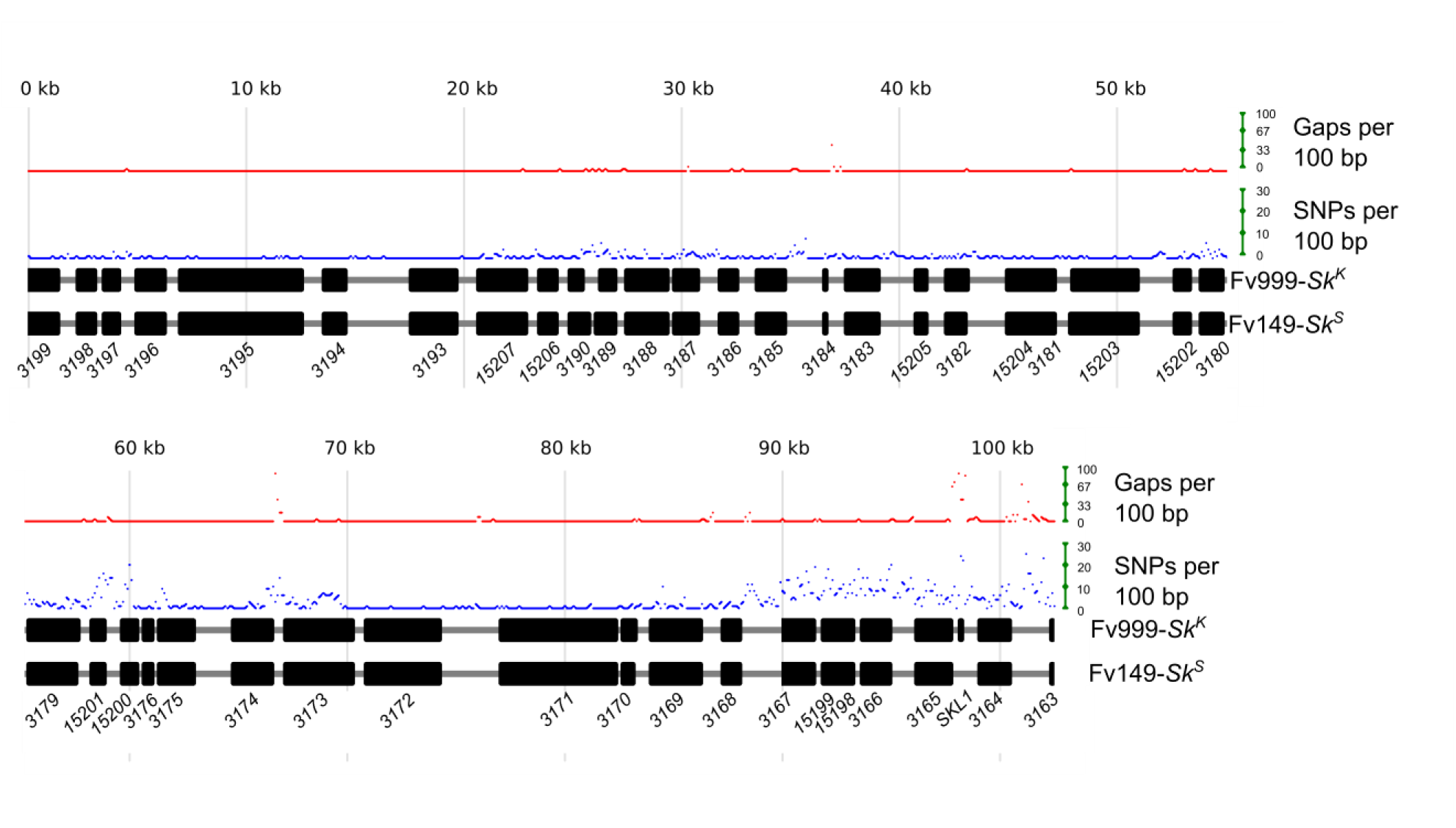
Strain Fv999-*Sk*^*K*^ carries a unique gene within a hypervariable region of the *Sk* locus. The *Sk* locus from Fv999-*Sk*^*K*^ (GenBank, KU963213) and Fv149-*Sk*^*S*^ (GenBank, CM000582.1, positions 665,669 to 767,411) were imported into BioEdit (Version 7.2.5) (Hall 1999) and aligned with ClustalW. Custom Perl scripts were used to examine base mismatches, gap positions, and to generate the diagram. The total number of mismatches (SNPs) between the two sequences was calculated for each 100-base, non-overlapping window of the alignment. The total number of gaps was also calculated for each 100-base, non-overlapping window. A gap position was not considered an SNP. By this definition, a window with 100 gaps can have no SNPs. Black rectangles represent the coding regions of predicted genes. Gene names are abbreviated according to their identification tags in the Fv149-*Sk*^*S*^ annotation. For example: *FVEG_03199*is shortened to *3199*. A striking feature of the alignment is the presence of at least three hypervariable regions; approximately spanning *FVEG_03180* to *FVEG_03175*, *FVEG_3174* to *FVEG_3173*, and *FVEG_03167*to the right border. A second striking feature is the presence of a unique gene, *SKL1*, in Fv999-*Sk*^*K*^ only.

The 102-kb *Sk* locus in strain Fv149-*Sk*^*S*^ includes 41 putative protein coding genes (Figure 4 and Table 4). The *Sk* locus in Fv999-*Sk*^*K*^ carries the same 41 genes as well as one additional protein coding gene, *SKL1* (for *Spore Killer Locus* 1), that is located between *FVEG_03165* and *FVEG_03164* (Figure 4). BLASTp analysis (Altschul *et al.* 1997) of the NCBI non-redundant protein database with the predicted 70-amino acid sequence of the putative Skl1 protein identified hypothetical proteins with Expect values ranging from 2 × 10^-49^ to 8 ×10^-04^ from various *formae speciales* of the *F. oxysporum* species complex. No significant hits were identified in other species (data not shown). A search of the NCBI Conserved Domain Database (Marchler-Bauer *et al.* 2010) failed to identify a domain within SKl1.

**Table 4.**
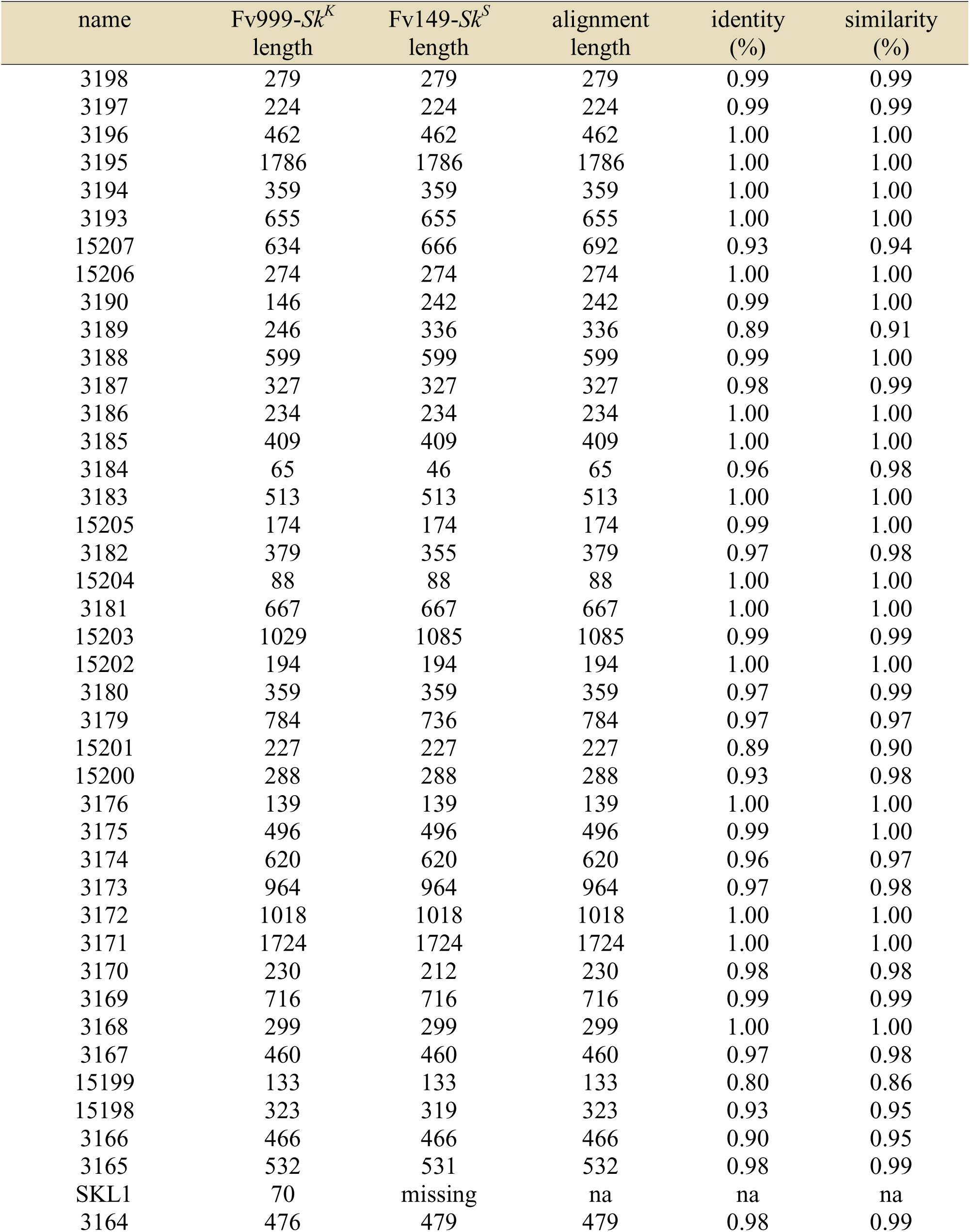
Comparison of predicted protein-coding sequences within the *Sk* locus of Fv999-*Sk*^*k*^ and Fv149-*Sk*^*S*^. The sequences of predicted proteins within the *Sk* locus of Fv999-*Sk*^*K*^ and Fv149-*Sk*^*S*^ were imported into Bioedit 7.2.5 (Hall 1999) and aligned with ClustalW (Thompson *et al.* 1994). Sequence alignments were then analyzed for percent identity and percent similarity. Gapped-positions were excluded from the identity and similarity calculations.

| name | Fv999-*Sk<sup>k</sup>*<br>length | Fv149-*Sk<sup>s</sup>*<br>length | alignment<br>length | identity<br>(%) | similarity<br>(%) |
| --- | --- | --- | --- | --- | --- |
| 3198 | 279 | 279 | 279 | 0.99 | 0.99 |
| 3197 | 224 | 224 | 224 | 0.99 | 0.99 |
| 3196 | 462 | 462 | 462 | 1.00 | 1.00 |
| 3195 | 1786 | 1786 | 1786 | 1.00 | 1.00 |
| 3194 | 359 | 359 | 359 | 1.00 | 1.00 |
| 3193 | 655 | 655 | 655 | 1.00 | 1.00 |
| 15207 | 634 | 666 | 692 | 0.93 | 0.94 |
| 15206 | 274 | 274 | 274 | 1.00 | 1.00 |
| 3190 | 146 | 242 | 242 | 0.99 | 1.00 |
| 3189 | 246 | 336 | 336 | 0.89 | 0.91 |
| 3188 | 599 | 599 | 599 | 0.99 | 1.00 |
| 3187 | 327 | 327 | 327 | 0.98 | 0.99 |
| 3186 | 234 | 234 | 234 | 1.00 | 1.00 |
| 3185 | 409 | 409 | 409 | 1.00 | 1.00 |
| 3184 | 65 | 46 | 65 | 0.96 | 0.98 |
| 3183 | 513 | 513 | 513 | 1.00 | 1.00 |
| 15205 | 174 | 174 | 174 | 0.99 | 1.00 |
| 3182 | 379 | 355 | 379 | 0.97 | 0.98 |
| 15204 | 88 | 88 | 88 | 1.00 | 1.00 |
| 3181 | 667 | 667 | 667 | 1.00 | 1.00 |
| 15203 | 1029 | 1085 | 1085 | 0.99 | 0.99 |
| 15202 | 194 | 194 | 194 | 1.00 | 1.00 |
| 3180 | 359 | 359 | 359 | 0.97 | 0.99 |
| 3179 | 784 | 736 | 784 | 0.97 | 0.97 |
| 15201 | 227 | 227 | 227 | 0.89 | 0.90 |
| 15200 | 288 | 288 | 288 | 0.93 | 0.98 |
| 3176 | 139 | 139 | 139 | 1.00 | 1.00 |
| 3175 | 496 | 496 | 496 | 0.99 | 1.00 |
| 3174 | 620 | 620 | 620 | 0.96 | 0.97 |
| 3173 | 964 | 964 | 964 | 0.97 | 0.98 |
| 3172 | 1018 | 1018 | 1018 | 1.00 | 1.00 |
| 3171 | 1724 | 1724 | 1724 | 1.00 | 1.00 |
| 3170 | 230 | 212 | 230 | 0.98 | 0.98 |
| 3169 | 716 | 716 | 716 | 0.99 | 0.99 |
| 3168 | 299 | 299 | 299 | 1.00 | 1.00 |
| 3167 | 460 | 460 | 460 | 0.97 | 0.98 |
| 15199 | 133 | 133 | 133 | 0.80 | 0.86 |
| 15198 | 323 | 319 | 323 | 0.93 | 0.95 |
| 3166 | 466 | 466 | 466 | 0.90 | 0.95 |
| 3165 | 532 | 531 | 532 | 0.98 | 0.99 |
| SKL1 | 70 | missing | na | na | na |
| 3164 | 476 | 479 | 479 | 0.98 | 0.99 |

To confirm that *SKL1* is absent in Fv149-*Sk*^*S*^, the sequences corresponding to the *FVEG_03165-FVEG_03164* intergenic region from Fv999-*Sk*^*K*^ and Fv149-*Sk*^*S*^ were aligned and examined. This intergenic region consists of 1,234 and 755 bases in Fv999-*Sk*^*K*^ and Fv149-*Sk*^*S*^ respectively (Figure S1). Examination of the alignment revealed that this difference in length was due to the presence of *SKL1* in Fv999-*Sk*^*K*^ and its absence in Fv149-*Sk*^*S*^ (Figure S1). To confirm that the absence of *SKL1* in Fv149-*Sk*^*S*^ was not due to an error in the reference genome sequence, we examined the lengths of the *FVEG_03165-FVEG_03164* intergenic region by PCR in our laboratory stocks of Fv999-*Sk*^*K*^ and Fv149-*Sk*^*S*^. The PCR product lengths were consistent with the presence of *SKL1* in Fv999-*Sk*^*K*^ and its absence in Fv149-*Sk*^*S*^ (Figure 5).

**Figure 5.**
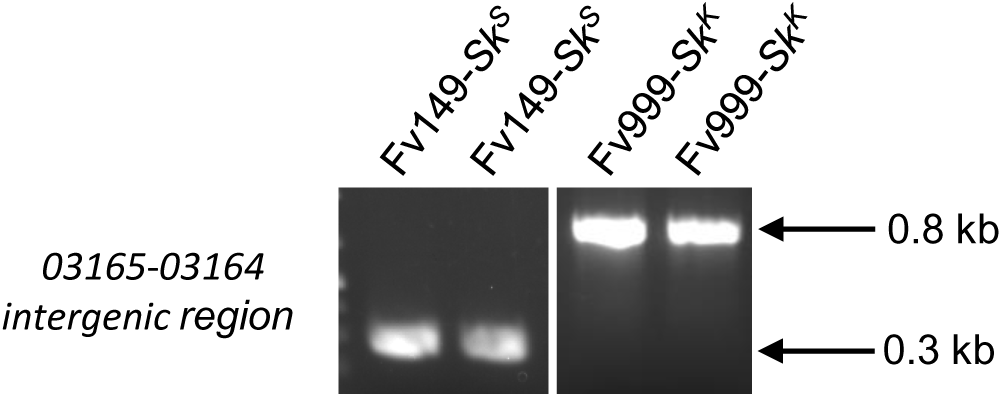
The *Sk* locus in Fv149-*Sk*^*S*^ is missing *SKL1*. To confirm that *SKL1* is missing from the *Sk* locus in Fv149-*Sk*^*S*^, genomic DNA was isolated from two liquid cultures of Fv149-*Sk*^*S*^ and two liquid cultures of Fv999-*Sk*^*K*^. All four genomic DNA samples were used as templates in standard PCR reactions with the following oligonucleotide primers: 5’ CGAATGACCTGGGGAGCCATAA 3’ and 5’ TCTCTCCACCACCTCCATCAGC 3’, which amplify the *FVEG_03165-FVEG_03164* intergenic regions in Fv999-*Sk*^*K*^ and Fv149-*Sk*^*S*^ PCR products were visualized by ethidium bromide staining after electrophoresis through a 1% agarose-TAE gel. We observed PCR products with lengths that are consistent with the absence of *SKL1* in Fv149-*Sk*^*S*^ (287 bp) and the presence of *SKL1* in Fv999-*Sk*^*K*^ (767 bp).

ClustalW alignments were used to investigate the level of polymorphism between proteins encoded by the *Sk* locus in Fv999-*Sk*^*K*^ and Fv149-*Sk*^*S*^ strains (Table 4). Analysis of these pairwise alignments revealed that Fveg_15199 is the most polymorphic protein of the locus. Only 80% of the 133 amino acids of Fveg_15199 are identical between the two strains. This is a remarkably high level of polymorphism given that 34 of the 41 shared proteins within the *Sk* locus of Fv999-*Sk*^*K*^ and Fv149-*Sk*^*S*^ are greater than 94% identical, and 40 of 41 are greater than 88% identical (Table 4). A BLASTp search of the NCBI non-redundant protein database identified homologs of Fveg_15199 in several *Fusarium, Neonectria, Neosartorya*, and *Aspergillus* species, among others (data not shown). A search of the NCBI Conserved Domain Database failed to identify a domain within Fveg_15199.

To shed light upon the transcriptional profile of protein-coding genes within the *Sk* locus, we analyzed five *F. verticillioides* RNAseq datasets from NCBI’s Sequence Read Archive. These datasets were for five time points following the induction of sexual development in a cross of Fv999-*Sk*^*K*^ × Fv149-*Sk*^*S*^ (Sikhakolli *et al.* 2012). Of the genes within the *Sk* locus, *FVEG_03171* exhibited the greatest increase in expression from the first to last time point (Table 5), suggesting that this gene may have an important role during later stages of sexual development. The corresponding protein, Fveg_03171, includes a WSC-domain (data not shown), which has been linked to carbohydrate-binding, cell wall integrity, and stress response. The gene *FVEG_03194* exhibited the greatest fold change over all five time points (>1050×) (Table 5). The Fveg_03194 protein does not have any recognized domains, although it is conserved in *Fusarium, Nectria, Trichoderma, Colletotrichum*, and a few other fungi (data not shown). Interestingly, *SKL1* reached the highest level of expression of all examined genes 2 hours after fertilization (Table 5). Furthermore, *SKL1* transcript levels were second only to *FVEG_03197* at the last time point (144 h, Table 5). The latter gene appears to be widely conserved among bacteria and eukaryotes (data not shown), but a function for it or its homologs is unknown.

**Table 5.** Expression analysis of genes from the *Sk* locus during sexual development. Sikhakolli *et al.* (2012) analyzed transcriptional changes in *F. verticillioides* during fruiting body development in Fv999-*Sk*^*K*^ × Fv149-*Sk*^*S*^ crosses by RNA sequencing (RNAseq) and deposited the datasets in NCBI’s Sequence Read Archive (Leinonen *et al.* 2011). The following datasets were downloaded from the database: 2 hours post fertilization (hpf) (SRR1592416), 24 hpf (SRR1592417), 48 hpf (SRR1592418), 72 hpf (SRR1592419), 96 hpf (SRR1592420), and 144 hpf (SRR1592421). Reads were aligned to coding sequences for the 42 genes (plus flanking genes *FVEG_03199* and *FVEG_03163*) from the *Sk* locus in Fv999-*Sk*^*K*^ with Bowtie 2 (Langmead and Salzberg 2012). RPKK values, or, reads per kilobase exon model per thousand mapped reads, were calculated for each gene. We then performed the same alignment and RPKK calculation procedure for the coding sequences of the 41 genes from the *Sk* locus (plus flanking genes *FVEG_03199* and *FVEG_03163*) in Fv149-*Sk*^*S*^. The RPKK values in the table are the average of the two RPKK calculations for each gene, except for *SKL1*, which is only found in Fv999-*Sk*^*K*^. RPKK is a variation upon RPKM as described by (Mortazavi *et al.* 2008). ^1^The minimum RPKK in the table. ^2^The maximum RPKK in the table. ^3^Fold change was calculated by dividing the maximum RPKK by the minimum RPKK to identify the genes undergoing the greatest expression change during the first 144 hours after fertilization. If the minimum RPKK for a gene was 0, it was arbitrarily assigned a minimum RPKK of 0.1 to approximate a fold change value.

| ID | 2 h | 24 h | 48 h | 72 h | 96 h | 144 h | Min<sup>1</sup> | Max<sup>2</sup> | Fold change<sup>3</sup> |
| --- | --- | --- | --- | --- | --- | --- | --- | --- | --- |
| 3198 | 39.17 | 150.92 | 38.40 | 24.98 | 28.10 | 20.10 | 20.10 | 150.92 | 7.51 |
| 3197 | 256.38 | 368.62 | 217.04 | 122.34 | 103.97 | 231.66 | 103.97 | 368.62 | 3.55 |
| 3196 | 0.83 | 0.13 | 0.87 | 0.56 | 0.83 | 0.87 | 0.13 | 0.87 | 6.59 |
| 3195 | 2.49 | 0.57 | 2.48 | 0.62 | 5.80 | 0.79 | 0.57 | 5.80 | 10.12 |
| 3194 | 0.25 | 0.12 | 0.30 | 14.02 | 124.53 | 1.20 | 0.12 | 124.53 | 1050.12 |
| 3193 | 138.42 | 94.82 | 71.69 | 43.35 | 80.19 | 75.41 | 43.35 | 138.42 | 3.19 |
| 15207 | 1.41 | 1.35 | 16.98 | 1.71 | 2.05 | 2.46 | 1.35 | 16.98 | 12.55 |
| 15206 | 0.78 | 1.40 | 5.23 | 1.29 | 0.76 | 1.08 | 0.76 | 5.23 | 6.86 |
| 3190 | 0.00 | 0.00 | 0.00 | 0.00 | 0.23 | 0.00 | 0.00 | 0.23 | 2.32 |
| 3189 | 0.00 | 0.00 | 0.00 | 0.05 | 0.50 | 0.00 | 0.00 | 0.50 | 4.95 |
| 3188 | 1.73 | 0.36 | 1.58 | 0.48 | 2.72 | 0.40 | 0.36 | 2.72 | 7.65 |
| 3187 | 1.90 | 0.97 | 1.34 | 31.36 | 42.16 | 0.71 | 0.71 | 42.16 | 59.47 |
| 3186 | 1.45 | 0.55 | 0.81 | 0.77 | 0.37 | 0.61 | 0.37 | 1.45 | 3.91 |
| 3185 | 5.41 | 3.74 | 6.17 | 2.61 | 1.72 | 1.51 | 1.51 | 6.17 | 4.09 |
| 3184 | 0.58 | 0.00 | 1.19 | 0.84 | 9.94 | 0.79 | 0.00 | 9.94 | 99.38 |
| 3183 | 59.62 | 22.63 | 32.99 | 13.12 | 29.15 | 21.10 | 13.12 | 59.62 | 4.54 |
| 15205 | 4.06 | 1.68 | 6.07 | 3.12 | 10.00 | 1.23 | 1.23 | 10.00 | 8.15 |
| 3182 | 0.30 | 0.00 | 1.19 | 0.00 | 0.29 | 0.00 | 0.00 | 1.19 | 11.93 |
| 15204 | 5.10 | 3.36 | 84.28 | 5.20 | 12.73 | 5.45 | 3.36 | 84.28 | 25.10 |
| 3181 | 2.16 | 0.50 | 5.24 | 0.50 | 1.93 | 0.66 | 0.50 | 5.24 | 10.57 |
| 15203 | 4.74 | 0.12 | 3.02 | 1.00 | 4.72 | 0.95 | 0.12 | 4.74 | 39.22 |
| 15202 | 1.05 | 0.44 | 8.61 | 7.61 | 9.45 | 0.74 | 0.44 | 9.45 | 21.57 |
| 3180 | 2.20 | 0.00 | 3.14 | 0.14 | 0.98 | 0.17 | 0.00 | 3.14 | 31.38 |
| 3179 | 1.28 | 0.06 | 1.14 | 0.10 | 2.70 | 0.09 | 0.06 | 2.70 | 47.85 |
| 15201 | 4.11 | 0.57 | 4.26 | 1.33 | 4.46 | 0.49 | 0.49 | 4.46 | 9.01 |
| 15200 | 1.02 | 0.55 | 0.32 | 0.36 | 0.29 | 0.24 | 0.24 | 1.02 | 4.16 |
| 3176 | 126.00 | 50.26 | 80.75 | 15.66 | 31.67 | 29.31 | 15.66 | 126.00 | 8.05 |
| 3175 | 48.87 | 12.93 | 38.03 | 9.13 | 19.60 | 13.41 | 9.13 | 48.87 | 5.35 |
| 3174 | 4.53 | 1.43 | 8.61 | 0.94 | 4.68 | 1.29 | 0.94 | 8.61 | 9.18 |
| 3173 | 1.04 | 0.17 | 2.86 | 0.42 | 1.90 | 0.42 | 0.17 | 2.86 | 17.29 |
| 3172 | 1.01 | 0.00 | 3.78 | 0.45 | 2.16 | 0.70 | 0.00 | 3.78 | 37.81 |
| 3171 | 2.95 | 37.34 | 38.55 | 117.07 | 44.37 | 94.58 | 2.95 | 117.07 | 39.66 |
| 3170 | 0.14 | 0.19 | 0.00 | 0.81 | 0.13 | 0.44 | 0.00 | 0.81 | 8.09 |
| 3169 | 5.48 | 0.61 | 5.38 | 3.02 | 7.40 | 2.61 | 0.61 | 7.40 | 12.05 |
| 3168 | 4.33 | 21.39 | 2.58 | 13.90 | 7.76 | 2.19 | 2.19 | 21.39 | 9.77 |
| 3167 | 0.17 | 0.71 | 0.13 | 0.21 | 0.12 | 0.04 | 0.04 | 0.71 | 19.47 |
| 15199 | 0.07 | 0.00 | 0.00 | 0.00 | 0.00 | 0.00 | 0.00 | 0.07 | 0.71 |
| 15198 | 1.34 | 0.12 | 0.56 | 0.21 | 0.31 | 0.19 | 0.12 | 1.34 | 11.68 |
| 3166 | 0.99 | 0.17 | 1.43 | 0.24 | 0.31 | 0.22 | 0.17 | 1.43 | 8.36 |
| 3165 | 20.77 | 11.51 | 13.49 | 9.68 | 8.41 | 4.09 | 4.09 | 20.77 | 5.07 |
| SKL1 | 481.58 | 392.59 | 227.42 | 211.66 | 360.52 | 97.65 | 97.65 | 481.58 | 4.93 |
| 3164 | 16.11 | 30.78 | 16.09 | 6.78 | 16.48 | 15.70 | 6.78 | 30.78 | 4.54 |

## DISCUSSION

The segregation of alternate alleles into separate gametes during meiosis encourages genetic conflict (Burt and Trivers 2008). Evidence for this is found in meiotic drive, which occurs when an allele is transmitted through meiosis in a biased manner. Meiotic drive elements are found in a diverse range of fungi, where they achieve biased transmission through sexual reproduction by killing meiospores carrying an alternate allele (Turner and Perkins 1979; Kathariou and Spieth 1982; Raju 1994; Dalstra *et al.* 2003; Grognet *et al.* 2014). Thus, fungal meiotic drive elements are often referred to as Spore killers. The molecular basis of meiotic drive by most Spore killers is unknown.

The existence of meiotic drive by spore killing in *Fusarium* was first recognized in 1982. More than three decades later, we still do not understand the molecular mechanism that mediates this process. This is similar to the situation in *Neurospora*, where three distinct spore killers, known as *Sk-1*, *Sk-2*, and *Sk-3*, were identified in 1979 (Turner and Perkins 1979). A breakthrough in *Sk-2* research was recently made by refining the location of an *Sk-2* resistance gene to a 52-kb sequence of DNA (Hammond, Rehard, Harris, *et al.* 2012). This provided the necessary foundation to clone and characterize the *Sk-2* resistance gene (Hammond, Rehard, Xiao, *et al.* 2012), which, in turn, allowed for the isolation of killer-less *Sk-2* mutants (Harvey *et al.* 2014). Here, we have taken a similar approach towards identifying the genetic basis for *Sk*^*K*^ in *F. verticillioides* by first refining the position of *Sk*^*K*^ to a 102-kb region of chromosome V.

As with previous work on *Sk*^*K*^ (Xu and Leslie 1996), we observed a biased transmission of *Sk*^*K*^-linked molecular markers (Figure 2B). This is not unexpected since the driving ability of *Sk*^*K*^ should also affect the transmission of alleles linked to *Sk*^*K*^. This phenomenon is referred to as genetic hitchhiking (Lyttle 1991). The closer a hitchhiker is to a meiotic driver, the more likely it is to be transmitted to the next generation through the sexual cycle. In our study, patterns of CAPS markers from the parent Fv999-*Sk*^*K*^ were inherited at a higher frequency than could be attributed to chance alone for five of six markers on chromosome V, and the transmission bias generally decreased with increasing distance from the *Sk* locus (Figure 2B). Surprisingly, CAPS-1 and CAPS-2 demonstrated higher levels of hitchhiking than CAPS-4 (Figure 2B), despite the latter being closer to the *Sk* locus (Figure 2D). One explanation for this could be the existence of a recombination hotspot between CAPS-4 and the *Sk* locus; however, we do not have additional data to support this hypothesis.

While a recombination hotspot could explain the unexpected decrease in hitchhiking for CAPS-4, the opposite phenomenon of recombination-suppression is often associated with meiotic drive elements (reviewed by Lyttle 1991). For example, *Neurospora Sk-2* requires specific alleles of at least two genes to mediate drive, a resistance gene called *rsk* and a killer gene called *rfk* (Hammond, Rehard, Xiao, *et al.* 2012; Harvey *et al.* 2014). Each gene is located on a different arm of chromosome III (Harvey *et al.* 2014). For *Sk-2* to succeed as a meiotic driver, it is imperative that an *Sk-2* ascospore inherit both *rsk* and *rfk* alleles because loss of *rsk* from *Sk-2* leads to a self-killing genotype (Hammond, Rehard, Xiao, *et al.* 2012). This helps explain why *Sk-2* is associated with a 30 cM long “recombination-blocked” region of chromosome III (Campbell and Turner 1987; Harvey *et al.* 2014). By suppressing recombination between *rsk* and *rfk*, *Sk-2* prevents these critical components of its meiotic drive mechanism from separating during meiosis. With respect to *F. verticillioides Sk*^*K*^, recombination suppression, if it exists at all, does not appear to be a significant phenomenon. For example, our CAPS marker analysis identified twelve recombination events between CAPS-3 and CAPS-4 (n = 59, Table S3), and two recombination events between CAPS-2 and CAPS-3 (n = 59, Table S3). This relatively high number of recombination events near the *Sk* locus argues against an *Sk-2*-like recombination block for *Sk*^*K*^.

If *F. verticillioides Sk*^*K*^ requires multiple genes to function as a meiotic drive element, all should be found within the 102-kb *Sk* locus defined by this study. For example, SNP-profiling of progeny JP98.75-*Sk*^*K*^, JP98.111-*Sk*^*K*^, and JP98.118-*Sk*^*K*^ indicated that only this region of chromosome V is common between them and the Fv999-*Sk*^*k*^ parent (Figures 2D and Figure 4). Assuming spore killing is mediated by two or more distinct genes within the *Sk* locus, the close proximity of these alleles may negate the requirement for a *Neurospora Sk-2*-like recombination block. Alternatively, meiotic drive and spore killing may be mediated by a single gene. A fungal precedent for this is seen in the het-s spore killing mechanism of *Podospora anserina* (Coustou *et al.* 1997; Saupe 2011). This system is controlled by two alternate alleles of a single gene, named *het-S* and *het-s*. The former, *het-s*, encodes the HET-s prion and is the meiotic driver, while the latter, *het-S*, encodes the HET-S protein. Interaction of the HET-s prion with HET-S causes HET-S to relocate to cell membranes, resulting in cell death, presumably through loss of membrane integrity (Seuring *et al.* 2012). The *het-s* ascospores escape cell death because they do not produce the HET-S protein, and thus the *het-s* prion is not toxic to them. This is one example of how meiotic drive and spore killing can be mediated by alternate alleles of a single gene in fungi. Grognet *et al.* (2014) have recently identified *spok1* and *spok2*, two additional spore killing genes in *P. anserina*, both of which appear to be single-gene-based meiotic drive systems.

In the current study, comparative sequence analysis revealed multiple differences between the 102-kb *Sk* locus from *Sk*^*K*^ and *Sk*^*S*^ strains of *F. verticillioides*. Presumably, one or more of these differences is the genetic basis for the different *Sk* phenotypes exhibited by the strains. An analysis of SNPs and gaps identified at least three hypervariable regions within the locus (Figure 4). The interspersed nature of these regions is somewhat surprising and suggests that they could be under the influence of different selective pressures than the less variable regions of the *Sk* locus. The most striking difference between the *Sk* locus is the presence the *SKL1* gene in *Sk^K^* strains and its absence in *Sk^S^* strains. Differences between the hypervariable regions and *SKL1* in *Sk^K^* versus *Sk^S^* strains raise at least two key questions: 1) are the hypervariable regions and/or *SKL1* responsible for spore killing, and 2) do they correspond to the *Sk^K^* allele that was previously defined by phenotypic analysis (Kathariou and Spieth 1982). The high levels of expression of *SKL1* throughout sexual development are consistent with its involvement in sexual reproduction. However, in addition to *SKL1* and the hypervariable regions, there are also numerous less dramatic sequence differences between genes and intergenic regions of the *Sk* locus in *Sk*^*K*^ and *Sk*^*S*^ strains. The current data does not rule out the possibility that one or more of these differences is responsible for the *Sk*^*K*^ and *Sk*^*S*^ phenotypes. As a result, our efforts are now focused on targeted deletions of *SKL1* and other regions of the *Sk* locus to determine the genetic basis of the spore killing phenotype in *F. verticillioides*.

## ACKNOWLEDGEMENTS

We thank the Fungal Genetics Stock Center (McCluskey *et al.*, 2010) for some of the strains used in this study and the Broad Institute of Harvard and MIT (www.broad.mit.edu) for access to the *Fusarium* Comparative Database. Dr. John Leslie (Kansas State University) and Dr. Frances Trail (Michigan State University) kindly donated Fv999-*Sk*^*K*^ and Fv149-*Sk*^*S*^ for this study. Chris McGovern and Nathane Orwig of NCAUR provided much appreciated technical assistance. We thank Dr. Sven Saupe (CNRS - Université de Bordeaux) for sharing his knowledge of the *het-s* spore killing mechanism. TMH was supported by a grant from the Eunice Kennedy Shriver National Institute of Child Health and Human Development of the National Institutes of Health (NICHD, 1R15HD076309-01). Mention of trade names or commercial products in this article is solely for the purpose of providing specific information and does not imply recommendation or endorsement by the US Department of Agriculture. USDA is an equal opportunity provider and employer.

## Figure and Table Legends

**Figure S1.**
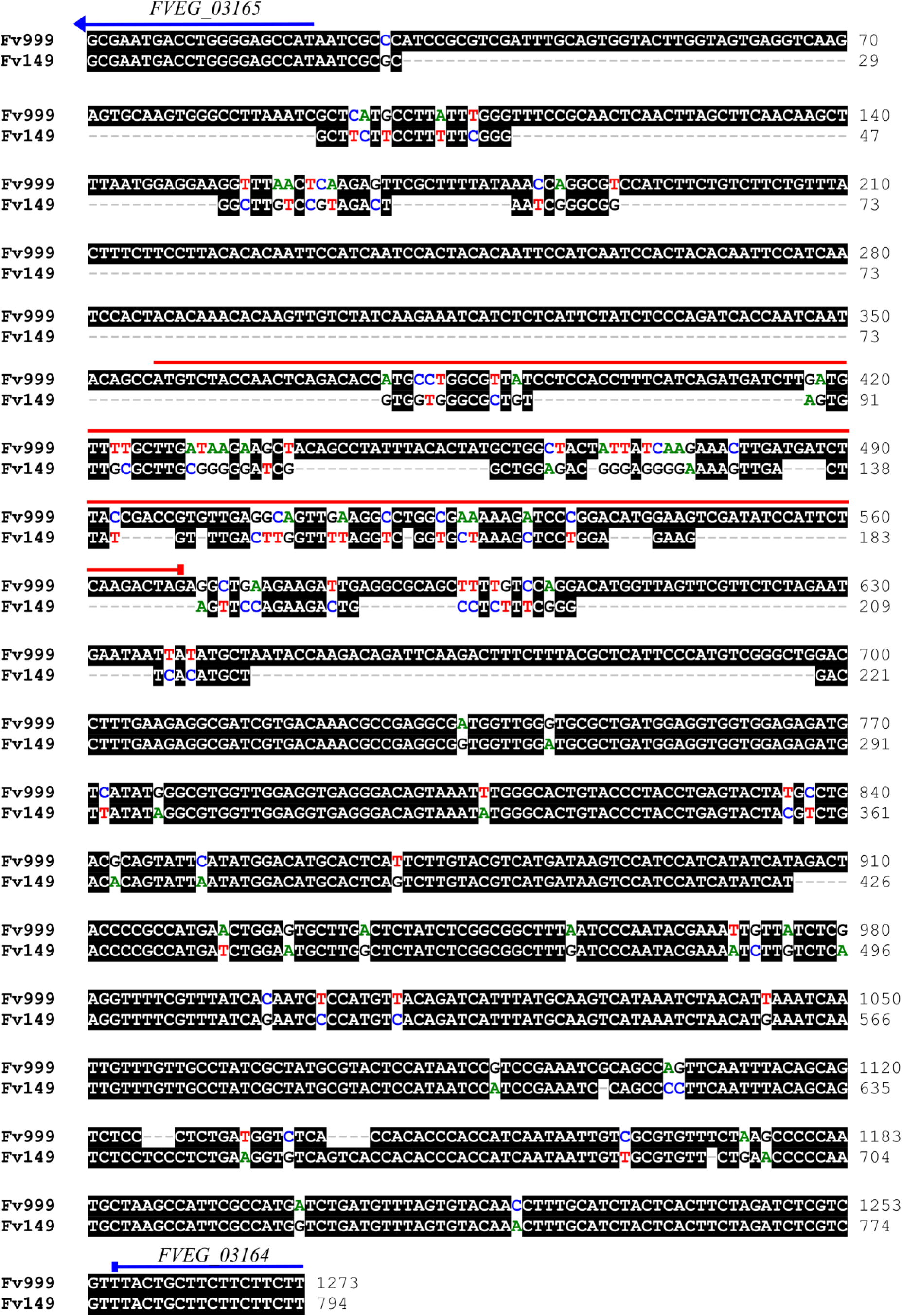
A ClustalW alignment of the *FVEG_03165-FVEG_03164* intergenic region from Fv999-*Sk*^*K*^ and Fv149-*Sk*^*S*^. The coding start site of flanking gene *FVEG_03165* is indicated with a blue arrow. Similarly, the coding stop site of flanking gene *FVEG_03164* is indicated with a blue line. The predicted coding sequence of *SKL1* is indicated with a red line. This alignment shows that *SKL1* is present in Fv999-*Sk*^*K*^ but not in Fv149-*Sk*^*S*^.

**Table S1.**
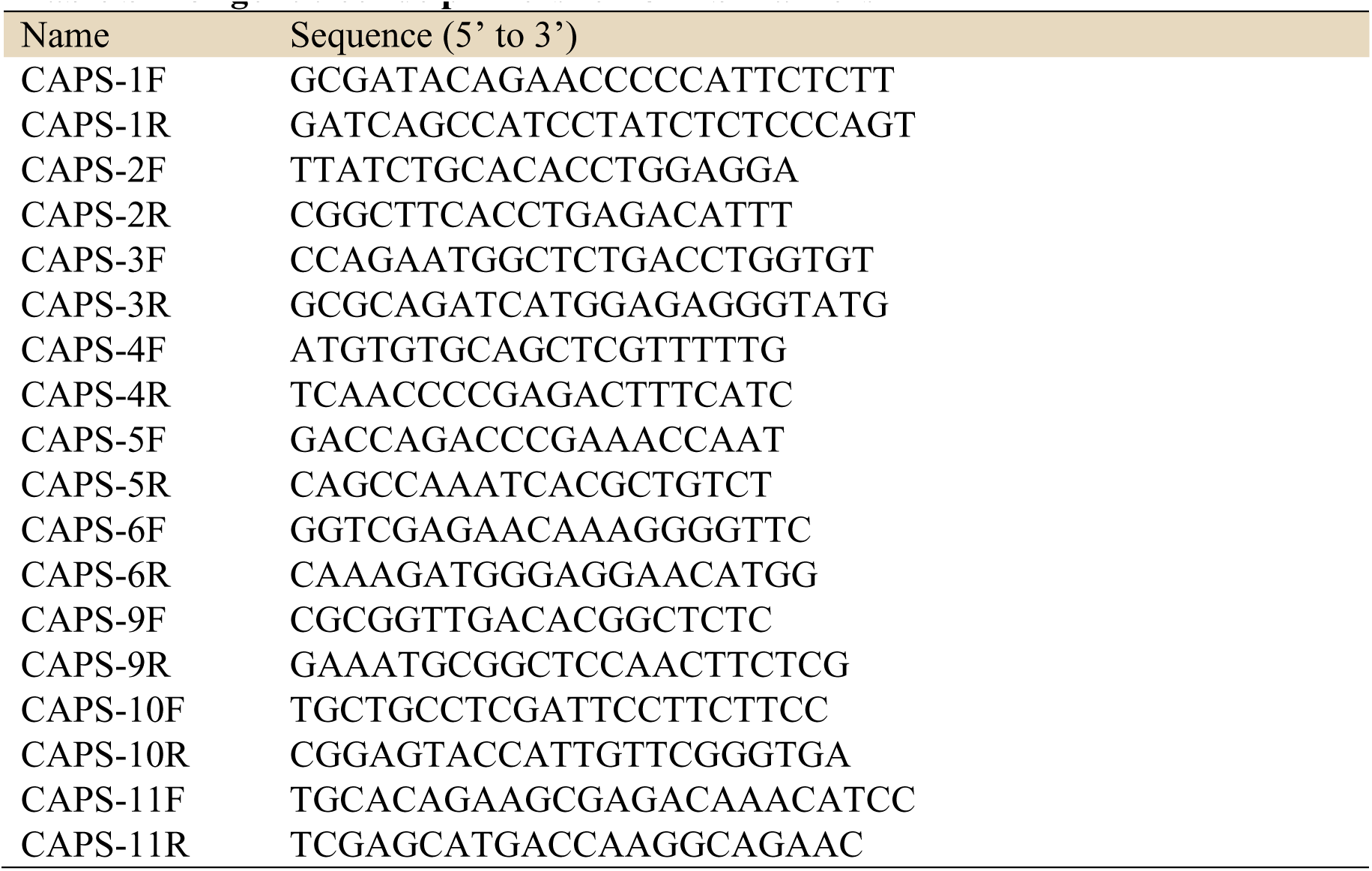
Oligonucleotide primers for CAPS markers

**Table S2.** CAPS marker sizes

| marker | PCR product length | Fv149 product<br>lengths after *Hae*III digest | Fv999 product<br>lengths after *Hae*III digest |
| --- | --- | --- | --- |
| CAPS-1 | 358 | 44, 86, 228, | 151, 207 |
| CAPS-2 | 307 | 132, 175 | 307 |
| CAPS-3 | 311 | 311 | 105, 206 |
| CAPS-4 | 288 | 110, 178 | 288 |
| CAPS-5 | 265 | 119, 146 | 265 |
| CAPS-6 | 325 | 34, 92, 199 | 34, 291 |
| CAPS-9 | 291 | 128, 163 | 291 |
| CAPS-10 | 404 | 404 | 68, 336 |
| CAPS-11 | 333 | 333 | 108, 225 |

**Table S3.** Genotypes of the Fv999-*Sk*^*K*^ × Fv149-*Sk*^*S*^ mapping population

| Strain | CAPS-5 | CAPS-4 | CAPS-3 | CAPS-2 | CAPS-1 | CAPS-6 | CAPS-9 | CAPS-10 | CAPS-11 |
| --- | --- | --- | --- | --- | --- | --- | --- | --- | --- |
| #60 | S | K | K | K | K | K | K | K | K |
| #61 | S | S | K | K | K | K | S | S | nd |
| #62 | K | K | K | K | K | K | S | K | S |
| #63 | K | K | K | K | K | K | K | K | S |
| #64 | K | K | K | K | K | K | S | K | K |
| #65 | K | K | K | K | K | K | S | K | S |
| #66 | K | K | K | K | K | K | K | K | K |
| #67 | S | K | K | K | K | K | S | S | S |
| #68 | K | K | K | K | K | K | S | S | nd |
| #69 | S | K | K | K | K | K | S | K | K |
| #70 | S | K | K | K | K | K | nd | S | S |
| #71 | S | S | K | K | K | K | nd | K | K |
| #72 | K | K | K | K | K | K | K | S | S |
| #73 | K | K | K | K | K | S | S | S | S |
| #74 | K | K | K | K | K | K | K | K | K |
| #75 | S | S | K | K | K | K | K | S | S |
| #76 | S | K | K | K | K | K | K | S | S |
| #77 | S | K | K | K | K | K | S | K | S |
| #78 | K | K | K | K | K | K | S | K | K |
| #79 | S | K | K | K | K | K | S | K | K |
| #80 | S | S | K | K | K | K | S | K | K |
| #81 | K | K | K | K | K | K | K | K | S |
| #82 | S | K | K | K | K | K | K | K | K |
| #83 | K | K | K | K | K | K | K | K | K |
| #84 | S | K | K | K | K | K | K | K | S |
| #85 | K | K | K | K | K | K | K | S | K |
| #86 | K | K | K | K | K | K | K | K | K |
| #87 | S | S | K | K | K | K | K | K | K |
| #88 | S | nd | K | K | K | K | K | S | S |
| #89 | S | K | K | K | K | K | K | S | K |
| #90 | S | K | K | K | K | K | K | K | S |
| #91 | K | K | K | K | K | K | K | K | K |
| #92 | S | K | K | K | K | K | K | nd | K |
| #93 | S | K | K | K | K | K | K | K | K |
| #94 | S | K | K | K | K | S | K | K | S |
| #95 | K | K | K | K | K | K | K | K | S |
| #96 | K | K | K | K | K | K | S | K | S |
| #97 | K | K | K | K | K | S | K | K | S |
| #98 | K | K | K | nd | K | K | K | K | K |
| #99 | K | K | K | K | K | K | K | S | S |
| #100 | S | K | K | K | K | K | K | K | S |
| #101 | S | S | K | K | K | K | K | S | K |
| #102 | K | K | nd | K | K | K | S | K | K |
| #103 | K | K | K | K | K | K | S | K | K |
| #104 | S | K | K | K | K | S | S | S | S |
| #105 | S | K | K | K | K | S | K | K | S |
| #106 | S | S | K | K | K | S | S | S | S |
| #107 | K | K | S | K | K | K | S | S | S |
| #108 | K | S | K | K | K | K | K | nd | K |
| #109 | K | K | K | K | K | K | K | K | S |
| #110 | S | S | K | K | K | S | S | S | K |
| #111 | K | S | S | S | S | S | K | S | K |
| #112 | K | K | K | K | K | K | S | nd | K |
| #113 | K | K | K | K | K | K | K | S | S |
| #114 | K | K | K | K | K | K | K | S | S |
| #115 | K | K | K | K | K | nd | S | S | K |
| #116 | S | S | K | K | K | K | K | K | K |
| #117 | K | K | K | K | K | K | K | S | K |
| #118 | K | K | K | S | S | S | K | K | K |
| #119 | S | S | K | K | K | K | K | S | K |

The chromosome V CAPS markers (1 through 6) are arranged in the table according to their relative order on the chromosome. K = Fv999-*Sk<sup>K</sup>* genotype; S = Fv149-*Sk<sup>S</sup>* genotype; nd = not determined.

